# Striatal cholinergic interneuron numbers are increased in a rodent model of dystonic cerebral palsy

**DOI:** 10.1101/2020.07.08.192922

**Authors:** Sushma Gandham, Yearam Tak, Bhooma R. Aravamuthan

## Abstract

Neonatal brain injury leading to cerebral palsy (CP) is the most common cause of childhood dystonia, a painful and functionally debilitating movement disorder. Rare monogenic etiologies of dystonia have been associated with striatal cholinergic interneuron (ChI) pathology. However it is unclear whether striatal ChI pathology is also associated with dystonia following neonatal brain injury. We used unbiased stereology to estimate striatal ChI and parvalbumin-positive GABAergic interneuron (PVI) numbers in a rodent model of neonatal brain injury that demonstrates electrophysiological markers of dystonia and spasticity. Striatal ChI numbers are increased following neonatal brain injury while PVI numbers are unchanged. These numbers do not correlate with electrophysiologic measures of dystonia severity. This suggests that striatal ChI pathology, though present, may not be the primary pathophysiologic contributor to dystonia following neonatal brain injury. Increased striatal ChI numbers could instead represent a passenger or protective phenomenon in the setting of dystonic CP.

## Introduction

Dystonia is a debilitating form of high muscle tone characterized by overflow muscle activation triggered by voluntary movement and worsened with heightened arousal (Albanese et al., 2013; Sanger et al., 2003). The most common cause of dystonia in childhood is neonatal brain injury resulting in cerebral palsy (CP), affecting at least 3 of every 10,000 live births (Ashwal et al., 2004; Himmelmann et al., 2009). Though deep grey matter injury, particularly of the striatum, has been implicated in dystonia pathogenesis in CP, gross striatal injury as visible on clinical imaging is neither necessary nor sufficient to yield dystonia following neonatal brain injury (reviewed by Aravamuthan and Waugh, 2016). Therefore the role of striatal injury in dystonia pathogenesis in CP remains unclear.

Dystonia pathogenesis has primarily been studied in rodent models of rare monogenic dystonia etiologies, all of which have demonstrated paradoxical striatal cholinergic interneuron (ChI) excitability (Eskow Jaunarajs et al., 2015). Striatal ChIs are decreased in number in at least one animal model of monogenic dystonia with striatal ChI loss occurring around the time of symptom onset (Pappas et al., 2015). Noting that dystonic CP differs from monogenic dystonias with regards to its clinical manifestation and treatment response, it is unclear whether dystonic CP is also associated with striatal ChI dysfunction.

Neonatal brain injury likely causes widespread effects on multiple different neuronal populations, with striatal injury implicated strongly in dystonic CP pathogenesis (Aravamuthan and Waugh, 2016). Therefore, any observed abnormalities in striatal ChIs in dystonic CP should be compared to abnormalities in other striatal interneurons to assess for specificity of striatal ChI abnormalities for dystonia. Parvalbumin-positive GABAergic striatal interneurons (PVIs) have not clearly been implicated in dystonia pathophysiology, but their inhibition has been associated with dyskinetic movements that resemble dystonia and they have been shown to either increase or decrease their firing rate time-locked to levodopa-induced dyskinesias (Alberico et al., 2017; Gittis et al., 2010). PVIs and ChIs also play unique and somewhat contrasting roles in movement regulation: PVI firing tends to precede movement and results in initiation of movement from rest while ChI firing occurs during ongoing movement and results in movement slowing or cessation (Gritton et al., 2019). Therefore, comparing ChI and PVI pathology in dystonic CP can be valuable to assess whether ChI-related abnormalities are specific to the dystonic phenotype.

We have recently developed a model of neonatal brain injury in rats that demonstrates electrophysiological characteristics of dystonia to varying degrees (Aravamuthan et al., 2020). This variability recapitulates clinical findings in that the presence and severity of dystonia in individuals with CP is variable, even in those who have undergone comparable mechanisms of neonatal brain injury (Lin et al., 2014). This animal model also demonstrates electrophysiological characteristics associated with spasticity, another distinct form of high tone seen more commonly in those with CP (Himmelmann et al., 2005). Spasticity is thought to be associated with injury to myelinated motor tracts, not deep grey matter as is typically associated with dystonia (Sanger et al., 2003). Here we examine striatal ChI and PVI number in rats following neonatal brain injury and the relationships between these interneuron numbers and electrophysiologic measures of dystonia and spasticity. Our aim is to determine whether striatal ChI dysfunction could be a contributing factor to dystonia following neonatal brain injury and whether the number of striatal ChIs or their distribution correlates with electrophysiologic measures of dystonia severity.

## Methods

### Animals

All procedures were approved by the Beth Israel Deaconess Medical Center Institutional Animal Care and Use Committee and the Washington University School of Medicine Institutional Animal Care and Use Committee. Timed pregnant Sprague Dawley dams were obtained from Charles River at embryonic days 17-18. Pups were born between embryonic days 21-23. Pups were aged within a 12-hour window from the time of birth and weaned at post-natal day (P) 21.

### Neonatal Brain Injury

Pups were either exposed to severe hypoxia at postnatal day (P) 7-8 or sham exposure, as has been previously described (Aravamuthan et al., 2020). Briefly, pups were placed in a hypoxia chamber (BioSpherix Ltd, Parish, NY) for 12 minutes with ambient oxygen concentrations decreasing from 21% to 0% fraction of inspired oxygen over that time. Approximately 6 minutes into the exposure, pups become pulseless, thus also contributing an ischemic element to this injury. P7-8 approximately corresponds to human term gestation (Clancy et al., 2007; Dobbing and Sands, 1979) which is of note because neonatal brain injury at term gestation may be clinically most likely to yield both dystonia and spasticity (Himmelmann et al., 2009).

### Electrophysiological assessments and signal processing

Between postnatal days 27-29, animals underwent nerve conduction and electromyographic (EMG) assessments to determine features that were analogous to dystonia and spasticity as has been previously described (Aravamuthan et al., 2020).

EMG quantification of co-contraction during voluntary hindlimb withdrawal movements was used as an analogue of dystonia. Dystonia is associated with overflow muscle activation triggered by voluntary movements and is exacerbated with increased arousal or stress. To capitalize on these clinical defining features, antagonist muscle co-contraction during voluntary isometric restrained hindlimb withdrawal movements were quantified using bipolar EMG recordings from the gastrocnemius/soleus complex and the tibilias anterior (Ambu Neuroline Subdermal Twisted Pair, Columbia, MD). Animals were deeply sedated with isoflurane (2 to 2.5% v/v in oxygen) and then had all four limbs and trunk restrained against the table surface using micro-pore tape. Animals were then allowed to emerge from anesthesia while inspiring 100% oxygen. Pressure to the base of the tail was applied with a bulldog clip every two seconds to elicit hindlimb withdrawal movements. EMG activity was recorded for 100 seconds beginning with the first voluntary hindlimb withdrawal movement in response to tail base pressure. The last 15 second epoch of this recording period (when animals were at the highest arousal state) was used for calculating co-contraction during voluntary hindlimb withdrawal attempts.

Digitized EMG signals were high-pass filtered at 65Hz and then rectified. EMG signal exceeding 5 standard deviations above the mean of 250 milliseconds of representative signal baseline was set at a value of 1 with the remainder of the signal set at 0 (Grosse et al., 2004). This leveling process preserves only the temporal information of motor unit firing while eliminating the confounding effects of muscle size and strength of contraction, which can vary between animals. Notably, the timing of motor unit firing is the only information needed to determine the presence of co-contraction between muscle pairs. Co-contraction was quantified using cross-correlogram analysis (*xcorr* function, MATLAB, MathWorks, Natick, MA). A higher cross correlogram peak amplitude at zero lag between EMG signals for the gastrocnemius/soleus and tibialis anterior is indicative of greater co-contraction between muscles. The cross-correlogram amplitude was divided by the total duration of movement in the analyzed epoch to allow comparison between animals who displayed differences in muscle contraction duration or frequency of hindlimb withdrawals (DeAndrade et al., 2016). The Fisher z-transform was then applied to yield a constant variance across records (DeAndrade et al., 2016).

The Hoffman reflex (H-reflex), the electrophysiological equivalent of the spinal stretch reflex, was measured as an analogue of spasticity. Spasticity is associated with hyperreflexia and spasticity severity is known to correlate with higher H-reflex amplitudes particularly with high frequency stimulation (Chen et al., 2001; Garrison et al., 2011; Hoving et al., 2006; Kumru et al., 2015; Pizzi et al., 2005; Ryu et al., 2017; Tekgül et al., 2013; Yates et al., 2008). H-reflexes were elicited in rats lightly sedated with isoflurane (0.5 to 1% v/v in oxygen) such that vibrissae and whisking reflexes were absent. The tibial nerve was stimulated using monopolar stimulating electrodes (Natus TECA electrodes, 1 inch length, 0.36 mm diameter) placed medial and lateral to the calcaneal tendon. The H-reflex response was recorded from the plantar interossei and occurred 5-8 msec after the stimulus (pulse width 0.2 msec, amplitude 0.5-3mA) (Yates et al., 2008). H-reflexes were elicited at 0.2 Hz, then at 2 Hz, then again at 0.2 Hz stimulation frequencies to examine H-reflex amplitudes and low and high frequency stimulation. At least five responses were elicited at each stimulus frequency to ensure H-reflex stability before recording five H-reflex responses for analysis. H-reflex suppression was measured as the ratio between the average H-reflex peak amplitude at 2 Hz divided by the average H-reflex peak amplitude at 0.2 Hz. H-reflex amplitude should decrease with 2 Hz stimulation (relative to 0.2 Hz stimulation) in healthy animals. That is, in healthy animals the H-reflex amplitude ratio should be less than 1. The H-reflex amplitude should be comparatively maintained at 2 Hz in animals with spasticity. That is, in spastic animals the H-reflex amplitude ratio should be close to 1.

### Tissue processing and immunohistochemistry

Rats were transcardially perfused with phosphate-buffered saline (PBS) and then 4% paraformaldehyde at P60. Brains were extracted and placed in 4% paraformaldehyde solution overnight. Brains were then transferred to 20% glycerol and 2% dimethylsulfoxide to prevent freeze-artifacts during slicing. The specimens were then embedded in a gelatin matrix using MultiBrain® Technology (NeuroScience Associates, Knoxville, TN).The specimens were rapidly frozen after curing by immersion in 2-Methylbutane chilled with crushed dry ice and mounted on a freezing stage of an American Optical 860 sliding microtome. Brains were sectioned coronally at 30 μm thickness. Sections were collected in Antigen Preserve solution (50% PBS pH7.0, 50% Ethylene Glycol, 1% polyvinyl pyrrolidone). No sections were discarded.

For immunohistochemistry, free floating sections were stained with choline-acetyl transferase (ChAT Millipore, Catalog Number AB144P, 1:3000 dilution) or parvalbumin (PVA, Swant, Catolog Number PV-235, 1:15000 dilution). Incubation solutions were in Tris buffered saline (TBS). Vehicle solutions contained Triton X-100 for permeabilization. Rinses were with TBS. After a hydrogen peroxide treatment and blocking serum, the sections were immunostained with the primary antibodies, and incubated overnight at room temperature. Following rinses, a biotinylated secondary antibody (anti-rabbit IgG for ChAT and anti-mouse IgG for PVA) was applied. After further rinses, avidin-biotin-HRP complex solution was applied (VECTASTAIN® Elite ABC, Vector, Burlingame, CA). The sections were again rinsed, then treated with diaminobenzidine tetrahydrochloride and hydrogen peroxide to create a visible reaction product. Following further rinses, the sections were mounted on gelatin coated glass slides and air dried. The slides were then dehydrated in alcohols, cleared in xylene, and coverslipped.

### Cell Counting

The number of ChIs and PVIs were estimated with unbiased stereology using the optical fractionator workflow in MBF Stereo Investigator (MBF Biosciences, Williston, VT) linked to a Zeiss Axio Imager Z2 Fluorescence Microscope with ApoTime 2 (Zeiss Microscopy, White Plains, NY). The dorsal striatum region of interest was first identified under 5x magnification. The medial border was defined by the lateral ventricle, the dorsal and lateral borders by the corpus callosum, and the ventral border by the anterior commissure. The dorsal striatum region of interest extended from 1.7 mm anterior to 0.3 mm posterior to bregma. Interneuron number estimation was performed at 40x magnification using an oil immersion lens. The stage controlling computer randomly placed the counting frame on the first counting area, and then systematically moved it until the entire selected field was sampled. A grid size of 300×300μm and counting frame of 150×150μm was used. Only the top of the cells that came into focus within the counting volume were counted. Guard zones of 2 μm at the top and bottom of each slice were used. Every sixth slice was assessed; that is 12 evenly distributed slices across the dorsal striatum were assessed with each slice 180 μm from the previously assessed slice. Striatal ChIs and PVIs were assessed for a subset of four brains twice to ensure consistent identification of cells. For all assessed specimens, the Gunderson coefficient of error was less than 0.05 with m=1 smoothing class. Counting was performed bilaterally with all data presented as the sum of both sides.

### Statistical Analyses

All statistical analyses were done with Graph Pad Prism 8 (GraphPad Software, San Diego, CA). Striatal ChI and PVI numbers, cross-correlogram amplitudes, and H-reflex amplitude ratios were compared between sham and hypoxia-exposed animals using t-tests. Linear regression analysis was used to examine the relationships between striatal ChI and PVI numbers and electrophysiologic measures of dystonia and spasticity. One-way ANOVAs with post-hoc Tukey HSD tests were used to compare interneuron numbers in sham-exposed mice to interneuron numbers in mice who displayed low (bottom 50%) and high (top 50%) dystonia and spasticity electrophysiologic measures.

## Results

### Striatal ChI numbers, but not PVI numbers, are increased in rats following neonatal brain injury

Striatal ChI numbers and PVI numbers in sham-exposed animals were comparable to what has been published previously (Larsson et al., 2001). In contrast to what has been observed in a mouse model of monogenic dystonia where striatal ChI numbers were decreased (Pappas et al., 2015), striatal ChI numbers were significantly increased in hypoxia-exposed animals (20564, 95% CI 19396-21732) compared to sham-exposed animals (15668, 95% CI 12373-18962). Striatal PVI number was not different between hypoxia-exposed animals (23951, 95% CI 21812-26089) and sham-exposed animals (24700, 95% CI 22659-26742) (Fig. 1).

**Figure 1.**
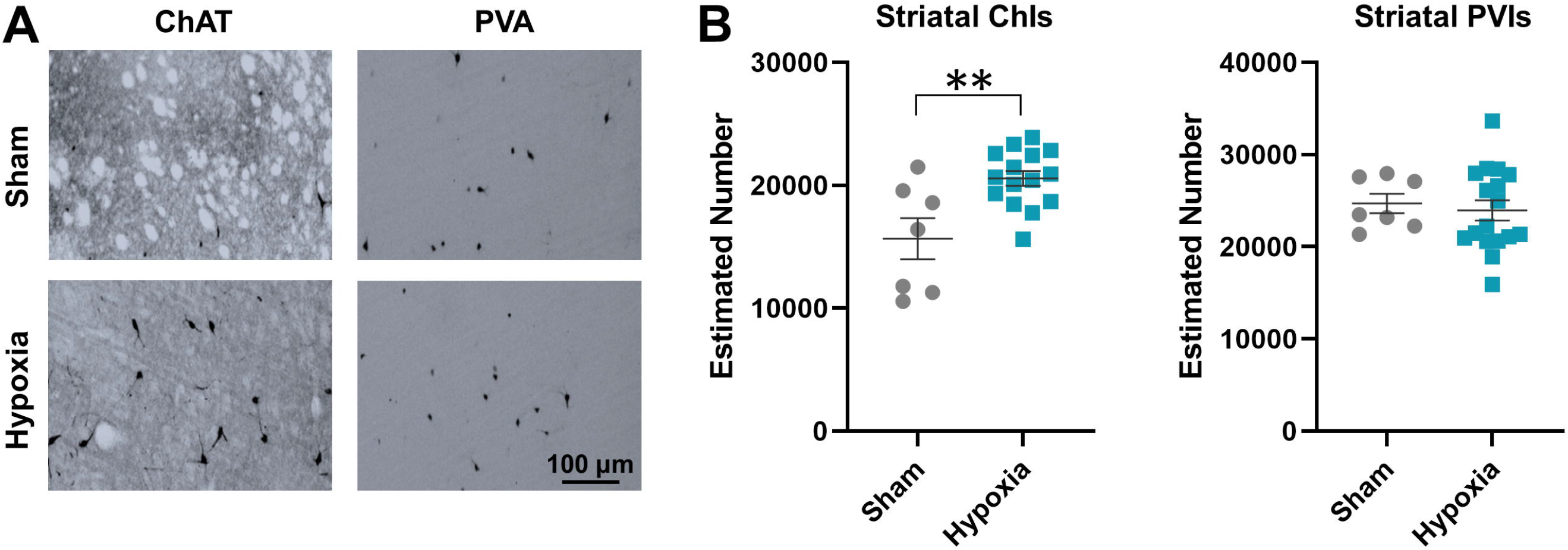
Striatal cholinergic interneuron (ChI) and parvalbumin-positive GABAergic interneuron (PVI) numbers in rats following neonatal hypoxia. A) Representative sections stained for choline-acetyl transferase (ChAT) and parvalbumin (PVA). B) Striatal ChIs, but not PVIs, are increased following neonatal hypoxia (Student’s t-test, **p<0.01). Error bars indicate mean +/− standard error of the mean.

### Striatal ChI and PVI numbers do not correlate with electrophysiologic measures of dystonia

Rats demonstrate significantly higher cross-correlogram amplitudes suggestive of dystonia following neonatal hypoxia exposure (0.8722 μV, 95% CI 0.7505-0.9939) compared to rats following neonatal sham exposure (0.5435 μV, 95% CI 0.4474-0.6395) (Fig. 2A). To examine whether there was a relationship between striatal interneuron numbers and cross-correlogram amplitude values, hypoxia-exposed animals were divided into two groups: those with cross-correlogram values in the bottom 50% of the group and those with values in the top 50% of the group. There was no difference between groups segregated by cross-correlogram values with regards to striatal ChI or striatal PVI numbers (one-way ANOVA followed by post-hoc Tukey HSD). Linear regression analysis also did not demonstrate any significant correlation between striatal ChI and PVI numbers and cross-correlogram amplitudes (Fig. 2B).

**Figure 2.**
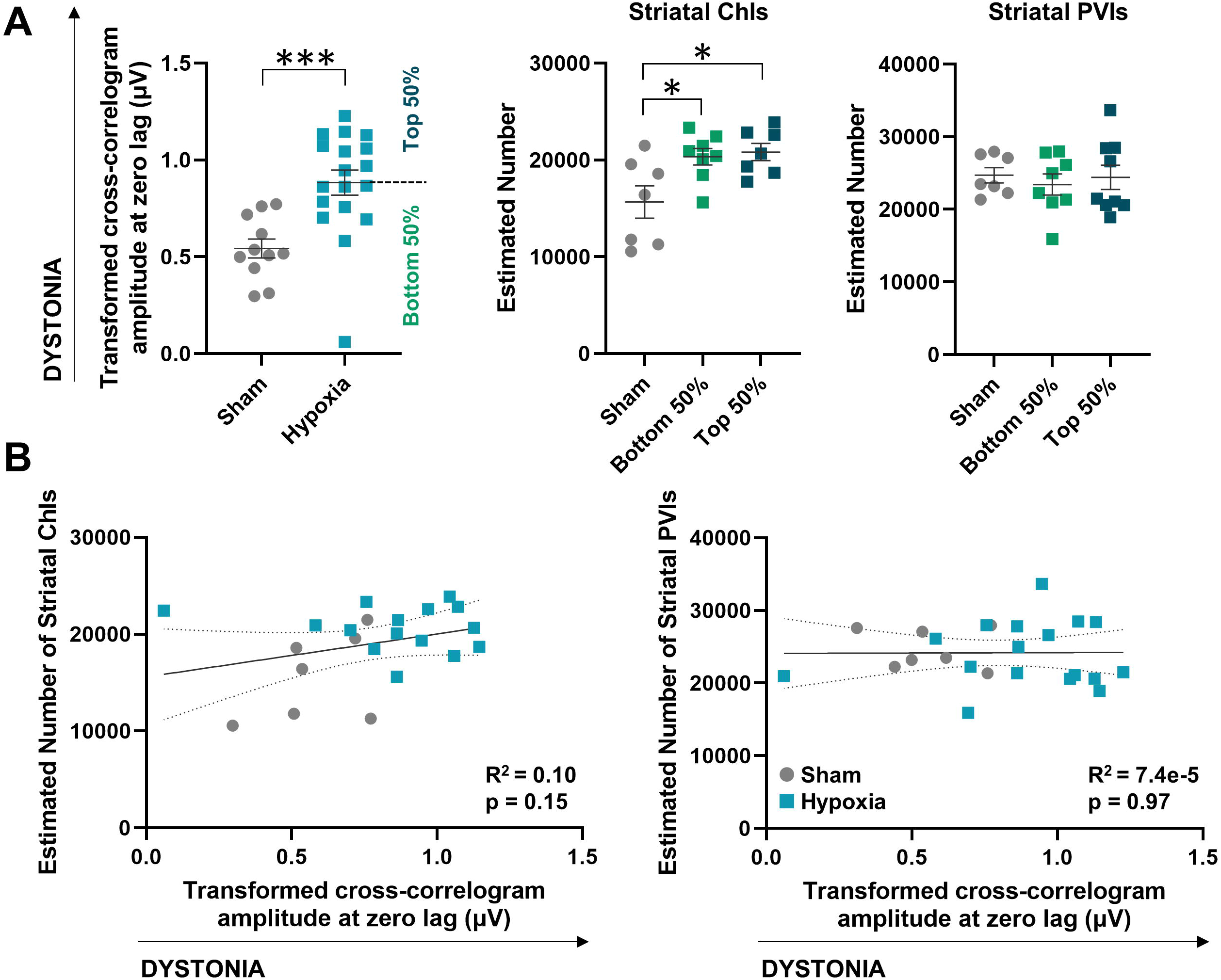
Relationship between striatal cholinergic interneuron (ChI) and parvalbumin-positive GABAergic interneuron (PVI) numbers and electrophysiologic measures of dystonia in rats following neonatal hypoxia. A) Arousal-provoked antagonist muscle co-contraction (cross-correlogram amplitude) is significantly greater in hypoxia-exposed animals compared to sham exposed animals (Student’s t-test, ***p<0.001). There is no difference between groups with low versus high cross-correlogram amplitudes with regards to striatal ChI or PVI numbers, though striatal ChIs remain significantly higher following hypoxia exposure compared to sham exposure regardless of associated cross-correlogram amplitude values (one-way ANOVA with post-hoc Tukey HSD, *p<0.05). Error bars indicate mean +/− standard error of the mean. B) Linear regression analyses reveal no correlation between cross-correlogram amplitudes and striatal ChI numbers (left) or PVI numbers (right). The linear regression line is solid. The 95% confidence band is dashed.

### Striatal ChI and PVI numbers correlate with electrophysiologic measures of spasticity

Rats following neonatal hypoxia exposure maintained H-reflex amplitudes with high frequency stimulation suggestive of hyperreflexia associated with spasticity (H-reflex amplitude ratio of 0.8667, 95% CI 0.5882-1.1453). In contrast, rats following neonatal sham exposure demonstrated H-reflex amplitude suppression with high frequency stimulation (H-reflex amplitude ratio of 0.3650, 95% CI 0.1899-0.5401) (Fig. 3A). To examine whether there was a relationship between striatal interneuron numbers and H-reflex amplitudes, hypoxia-exposed animals were divided into two groups based on H-reflex amplitude ratios: those with ratios in the bottom 50% of the group (thought to be less spastic) and those with ratios in the top 50% of the group (thought to be more spastic). There was no difference between these groups with regards to striatal ChI or striatal PVI numbers (one-way ANOVA followed by post-hoc Tukey HSD). Surprisingly, noting that deep grey matter injury is not pathophysiologically linked to spasticity, linear regression analysis revealed a weak positive correlation between striatal ChI number and H-reflex ratios, suggesting that larger striatal ChI numbers might be found in animals demonstrating higher spasticity measures. There was also a significant negative correlation between striatal PVI numbers and H-reflex ratios, suggesting that fewer striatal PVIs may be found in animals demonstrating higher spasticity measures (Fig. 3B).

**Figure 3.**
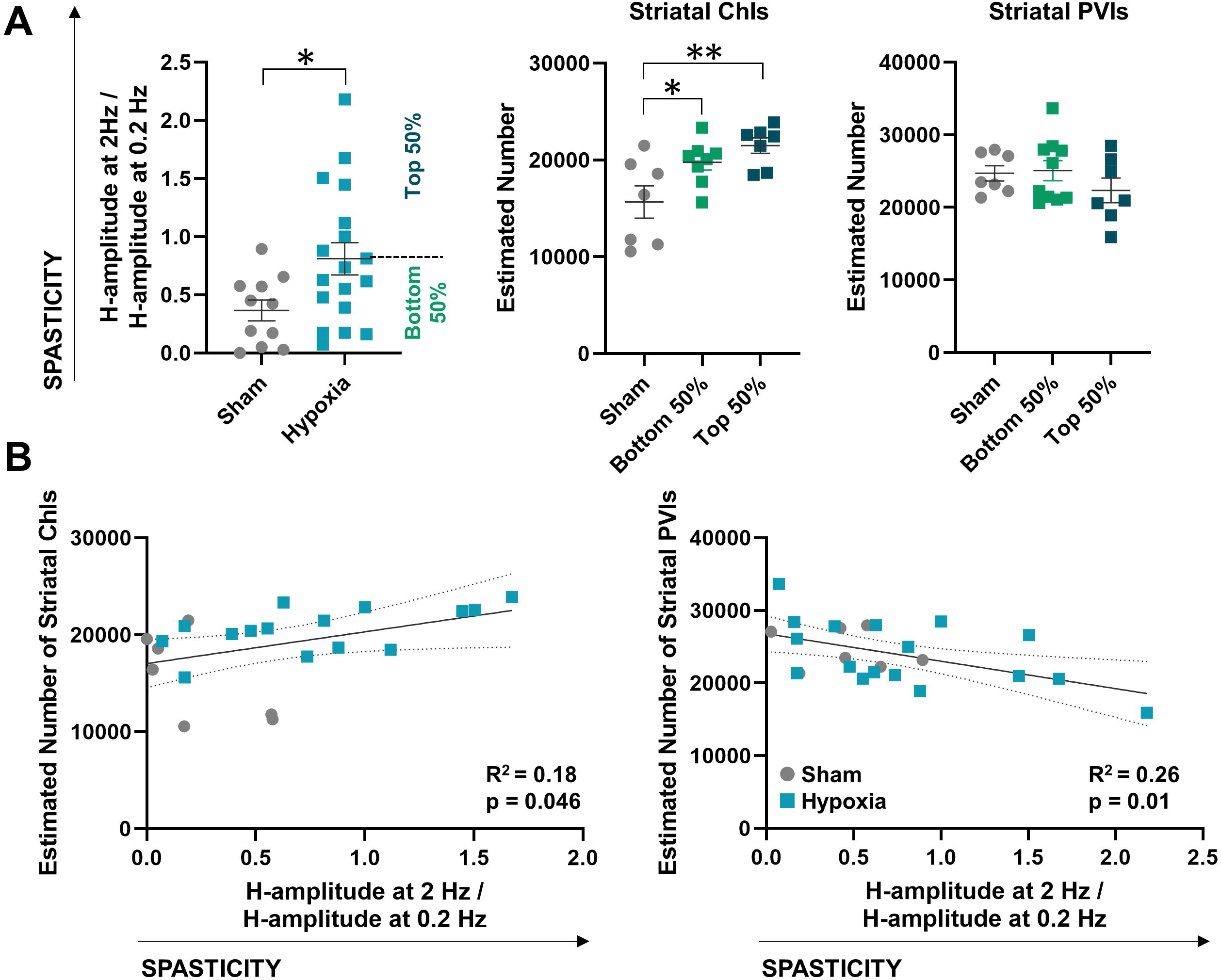
Relationship between striatal cholinergic interneuron (ChI) and parvalbumin-positive GABAergic interneuron (PVI) numbers and electrophysiologic measures of spasticity in rats following neonatal hypoxia. A) The ratio of H-reflex amplitude between high and low frequency stimulation is significantly greater in hypoxia-exposed animals compared to sham-exposed animals (Student’s t-test, *p<0.05). There is no difference between groups with low versus high H-reflex amplitude ratios with regards to striatal ChI or PVI numbers, though striatal ChIs remain significantly higher following hypoxia exposure compared to sham exposure regardless of associated H-reflex amplitude values (one-way ANOVA with post-hoc Tukey HSD, *p<0.05, **p<0.01). Error bars indicate mean +/− standard error of the mean. B) Linear regression analyses reveal weak but significant correlations between H-reflex amplitude ratios and striatal ChI numbers (left) and PVI numbers (right). The linear regression line is solid. The 95% confidence band is dashed.

## Discussion

This study demonstrates that neonatal brain injury in rats results in increased striatal ChI number regardless of whether the rat displays high or low electrophysiological markers for dystonia. PVI number is unaffected following neonatal brain injury in rats. Unexpectedly, striatal ChI and PVI numbers appear to weakly correlate with an electrophysiologic analogue of spasticity.

Increased striatal ChI number in this rat model of dystonic cerebral palsy regardless of the apparent electrophysiological severity of dystonia suggests that changes in striatal ChI number may not directly contribute to dystonia pathogenesis following neonatal brain injury. However, the fact that striatal ChIs are increased following a severe neonatal hypoxic event is notable. This finding could represent a protective phenomenon particularly given that reduced striatal ChI number occurs concurrently with dystonic symptom onset in a rodent model of a monogenic dystonia (Pappas et al., 2015). It is additionally notable that striatal PVIs, a key GABAergic interneuron population important for motor control, are unchanged in number following neonatal brain injury. Therefore, the observed increase in striatal ChIs may be specific to this particular striatal interneuron population.

The mechanism of increased striatal ChIs following neonatal brain injury is unclear. As striatal neurogenesis is complete by P3 and peaks between embryonic day 15-16 (Marchand and Lajoie, 1986), it is unlikely that neonatal brain injury at P7 results in differentiation of other striatal interneurons into striatal ChIs. The observed increase may instead be due to increased striatal ChI ChAT expression as has been previously hypothesized (Larsson et al., 2001). That is, striatal ChI numbers may not actually be increased, but striatal ChI acetylcholine production might be. This would be in line with findings in some rodent models of monogenic dystonias which demonstrate increased extracellular acetylcholine levels (Eskow Jaunarajs et al., 2015).

The association between striatal ChI and PVI numbers and H-reflex amplitude ratios are difficult to interpret as directly linked phenomena. Though striatal electrical stimulation has historically been shown to modulate the amplitude of spinal reflexes, this has not been robustly replicated and there is no clarity regarding whether striatal electrical stimulation may have inadvertently involved the internal capsule or other corticospinal tracts (Segundo et al., 1958). Therefore, there remains no clear anatomical mechanism for striatal pathology to affect spinal stretch reflexes. This finding may instead reflect how striatal pathology following neonatal brain injury may co-occur with injury to corticospinal tracts as is often observed clinically (Pierson et al., 2007).

Future studies can assess the degree of corticospinal tract injury in this model and determine whether that correlates with measures of spasticity. Further histological characterization of other striatal and deep grey matter neuronal populations in this model may reveal associations with dystonia measures that were not apparent here. Finally, to more directly address the question of whether striatal ChIs play a causative role in dystonia pathogenesis, electrophysiological measures of dystonia can be assessed following excitation of striatal ChIs.

In sum, these results show that striatal ChI pathology is seen in a rodent model of dystonic cerebral palsy. The observed increase in striatal ChI number is in contrast to the observed decrease in striatal ChI number seen in a rodent model of monogenic dystonia (Pappas et al., 2015). However, it remains unclear whether striatal ChI pathology is causative of a dystonic phenotype following neonatal brain injury.

## Author Roles

B.R.A was responsible for the conception of the study. S.G. and B.R.A were responsible for initial drafting of the manuscript and figures. S.G., Y.T, and B.R.A contributed to the design of the study, data acquisition, and data analysis.

